# Preparation of single-cell suspension from mouse breast cancer focusing on preservation of original cell state information and cell type composition

**DOI:** 10.1101/824714

**Authors:** Abaffy Pavel, Lettlova Sandra, Truksa Jaroslav, Kubista Mikael, Sindelka Radek

## Abstract

Single-cell analysis of gene expression has become a very popular method during the last decade. Unfortunately, appropriate standardization and workflow optimization remain elusive. The first step of the single cell analysis requires that the solid tissue be disassociated into a suspension of individual cells. However, during this step several technical bias can arise which can later result in the misinterpretation of the data. The goal of this study was to identify and quantify the effect of these technical factors on the quality of the single-cell suspension and the subsequent interpretation of the produced expression data. We tested the effects of various enzymes used for dissociation, several centrifugation forces, dissociation temperatures and the addition of Actinomycin D, a gene expression inhibitor. RT-qPCR was used to assess the effect from each parameter alteration, while a single-cell RNA sequencing experiment was used to confirm the optimized factors. Our concluding results provide a complete protocol for the tissue dissociation of mouse mammary tumour from 4T1 cells that preserves the original cell state and is suitable for any single-cell RNA sequencing analysis. Furthermore, our workflow may serve as a guide for the optimization of the dissociation procedure of any other tissue of interest, which would ultimately improve the reproducibility of the reported data.

## INTRODUCTION

It has been more than 10 years since Tang *et al*. (2009) published the first paper on the analysis of the whole transcriptome from single cells (single-cell RNA-Seq, scRNA-Seq). Since then, scientists have produced optimized protocols for the preparation of sequencing libraries from individual cells. Such protocols have allowed for many different discoveries from the cell such as gene expression, chromatin modifications, copy number variation, and other “omics” (such as proteomics) (Guo et al., 2017; Cheung et al., 2018; Hou et al., 2016; Andor et al., 2018). However, the first step utilized by all these protocols is the preparation of the single-cell suspension, also known as tissue dissociation. Unfortunately, cells react to the stress induced by the disassociation process and this often leads to an artificial induction of gene expression. The level of this artificial induction is dependent on the particular dissociation protocol and sample type (van den Brink et al., 2017; Adam, Potter and Potter, 2017; Wu et al., 2017) and is confounded within the original expression pattern which can lead to biological misinterpretations (van den Brink et al., 2017). Recently, the problem of dissociation associated induced expression was discussed and suggested as a problematic factor during single cell analyses (van den Brink et al., 2017). During the dissociation process, certain genes within the cells have a sudden burst in expression. These genes usually consist of the stress induced or immediate early genes (IEGs). Interestingly, this set of genes is conserved in higher eukaryotes and has been observed after injury in mouse (Grose et al., 2002), fish (Ishida et al., 2010) and also during embryonic wound healing at the gastrula and tailbud stages in *Xenopus* embryos (Ding et al., 2017; Abaffy et al., in press). Usually the peak expression of these genes appears between 30 and 60 minutes after treatment/injury. Additionally, upregulation of many IEGs has also been observed in post-mortem tissue derived from mouse and fish (Pozhitkov et al., 2017). This particular group of genes consists of members within the AP1 pathway and heat-shock proteins.

One of the suggested solution for this issue is to filter out cells expressing IEGs during data analysis (van den Brink et al., 2017). Unfortunately, this approach would also eliminate cells that were naturally responding to other innate biological stress situations. Another solution is to modify the dissociation protocol to prevent/reduce induction of the IEGs expression. The first approach from this solution would be to perform dissociation and all subsequent steps at a lower temperature, which would significantly reduce the metabolic processes within the cells (Adam, Potter and Potter, 2017). However, a drawback of this approach is that the lower temperature would also decreases the efficacy of the dissociation enzymes. This would result in an incomplete dissociation of the tissue and the subsequent loss of some cell populations. The second approach is to use a transcription inhibitor to reduce IEGs expression during dissociation. Addition of Actinomycin D (ActD) has alreadu been successfully used during dissociation of neuronal tissue into single-cell suspension, where it was found to greatly reduced IEGs expression (Wu et al., 2017). However, the routine usage of inhibitors is limited because of their concentration and cell type dependent toxicity. The third approach is to use fixed tissue (Machado et al., 2017), which would render biological processes inactive. However, this approach requires the fixation of the tissue in paraformaldehyde (PFA) which leads to the cross-linking of proteins with RNA and can lead to a worse quality of the RNA. Interestingly, Machado *et al*. (2017) found no alteration in mRNA yield, quality, and composition when using fixed samples. However, they only analysed the effect of PFA fixation at the bulk level.

The last approach is single-nucleus RNA sequencing (snRNA-Seq) using flash-frozen tissue (Habib et al., 2016; Habib et al., 2017; Krishnaswami, S. R. et al., 2016; Lacar, B. et al., 2016). We *et al*. (2019) observed that there was no artificial transcriptional stress response when using snRNA-Seq. However, an important question remains on whether the nuclear RNA content represents well the whole cell transcriptome. Comparative studies (Grindberg, R. V. et al., 2013; Gao, R. et al., 2017; Habib et al., 2017) have shown an overall high correlation between scRNA-Seq and snRNA-Seq. However, small variations between the two methods have been observed in the content of poly-adenlyated ncRNAs (Krishnaswami, S. R. et al., 2016), enrichment of lncRNAs (Gao, R. et al., 2017) and abundance of intronic sequences (Grindberg, R. V. et al., 2013; Gao, R. et al., 2017; Habib et al., 2017). These biases must be removed during data analysis.

An additional obstacle during tissue dissociation is the reduction or even loss of cell subpopulations. Contrarily to the previously described problems, the impact of this loss on the interpretation of the final data is not sufficiently discussed in the literature. Recent studies have already touched on some of these issues that may arise during tissue dissociation (Potter and Potter, 2019). These studies have warned that scientists should not expect an accurate representation of the cell-types from data derived from the scRNA-Seq. A general suggestion to improve the quality of the single-cell suspension is through protocol optimization. However this is necessary for each type of tissue as there is no “one correct solution” for all types (Vieira Braga and Miragaia, 2019; Potter and Potter, 2019).

Tumour heterogeneity has been described many times in the literature (Winterhoff et al., 2017; Tirosh et al., 2016; Puram et al., 2017; Patel et al., 2014; Hou et al., 2016) and large-scale single-cell analyses performed from tumour biopsies or whole tumour tissues represent a great challenge. In addition, the complexity of a single tumour cell analysis comes from combined regulation by both intertumoural and extratumoural factors (Lawson et al., 2018). The critical aspect of these analyses is preservation of the original intact tumour signatures in terms of its cell populations, types and states (Lawson et al., 2018). Our goal here is to demonstrate questions and solutions during protocol optimization for single-cell suspension preparation with the main challenge of reduction of IEGs expression. Our work discusses the effect of the different protocol steps and their impact on the results interpretation.

## RESULTS

The process of preparing the single-cell suspension from the fresh solid tissue includes several modifiable steps (**Figure 1**). We utilized a 10x Genomics protocol as a gold standard for single-cell suspension preparation (10x Genomics, 2018; 10x Genomics, 2017). The protocol included: lysis of the erythrocytes to prevent issues with counting of the viable cells when using an automatic cell counter; removal of debris which would otherwise contribute to a high RNA background; and removal of dead cells which would reduce the cell viability. We optimized this protocol by using a low dissociation temperature (4 °C) as well as with the addition of ActD during dissociation, so as to minimize the dissociation-associated stress response. We tested also different enzymes, their concentrations and the effect of centrifugation on cell suspension quality.

**Figure 1:**
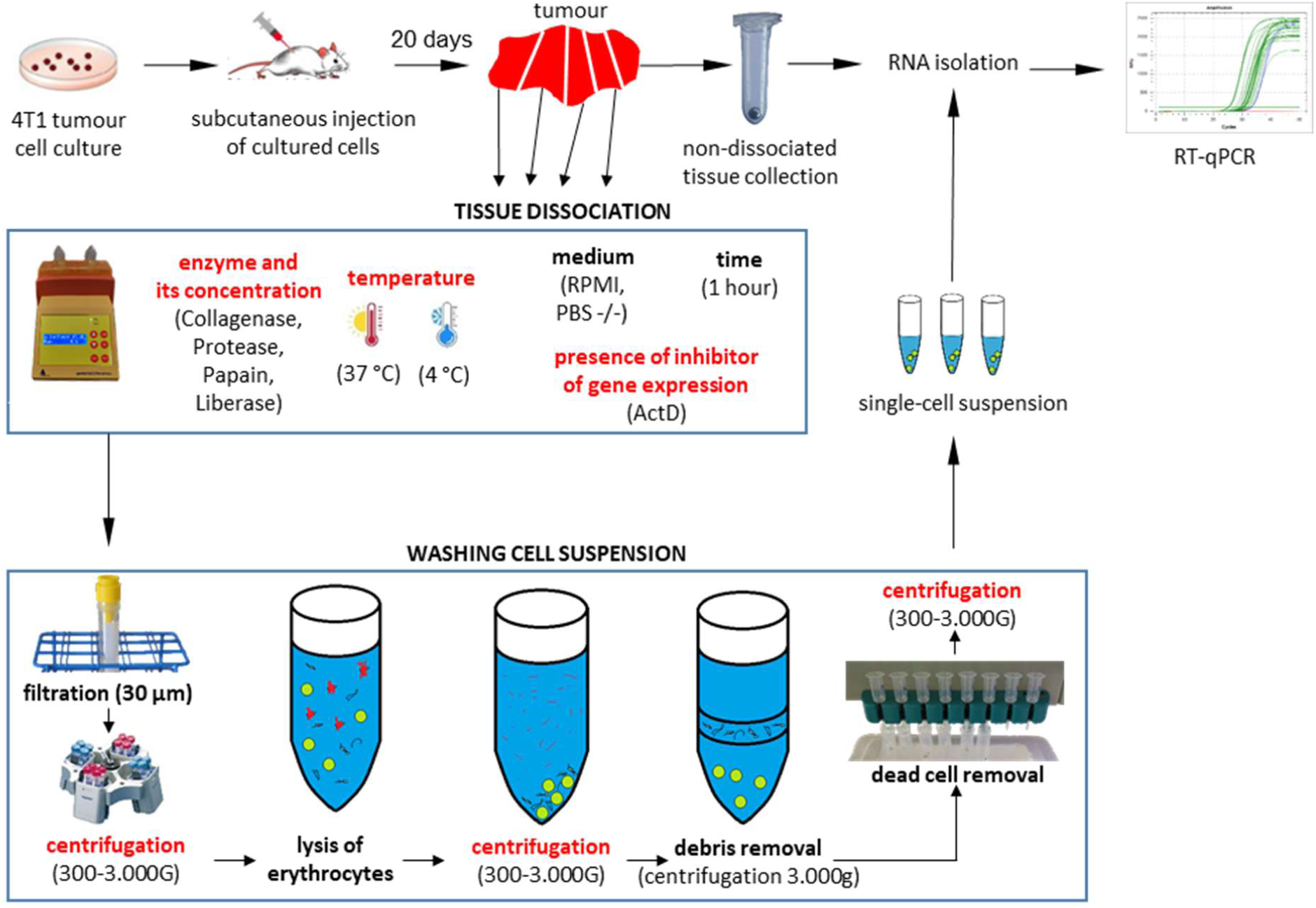
Scheme of the process of preparation of the single-cell suspension from the tumour tissue. Factors, which were optimized, are marked by red colour.

### Selection of dissociation enzyme

In the first step of the protocol optimization, we tested various dissociation enzymes using the recommended parameters. We also altered the following factors that may also contribute to the artificial gene expression via stimulation/degradation: i) low dissociation temperature (4 °C); ii) presence of transcription inhibitor ActD; iii) centrifugation at 300 g. The specific enzymes and their concentrations were selected based on available commercial and scientific protocols (in total ten different conditions, (**Figure 2A**): Collagenase, which was used in previous experiments during dissociation at 37 °C; Protease, which was used by Adam, *et al*. (2017); Papain, which is often used for dissociation of embryonic tissues at low temperature; Liberase™, a mixture of enzymes with expected higher activity in comparison to Collagenase. The Ca^2+^ ions were added into medium, because it is important for activation of Collagenase. Negative control was performed as a mechanical disruption in the absence of any enzyme.

**Figure 2:**
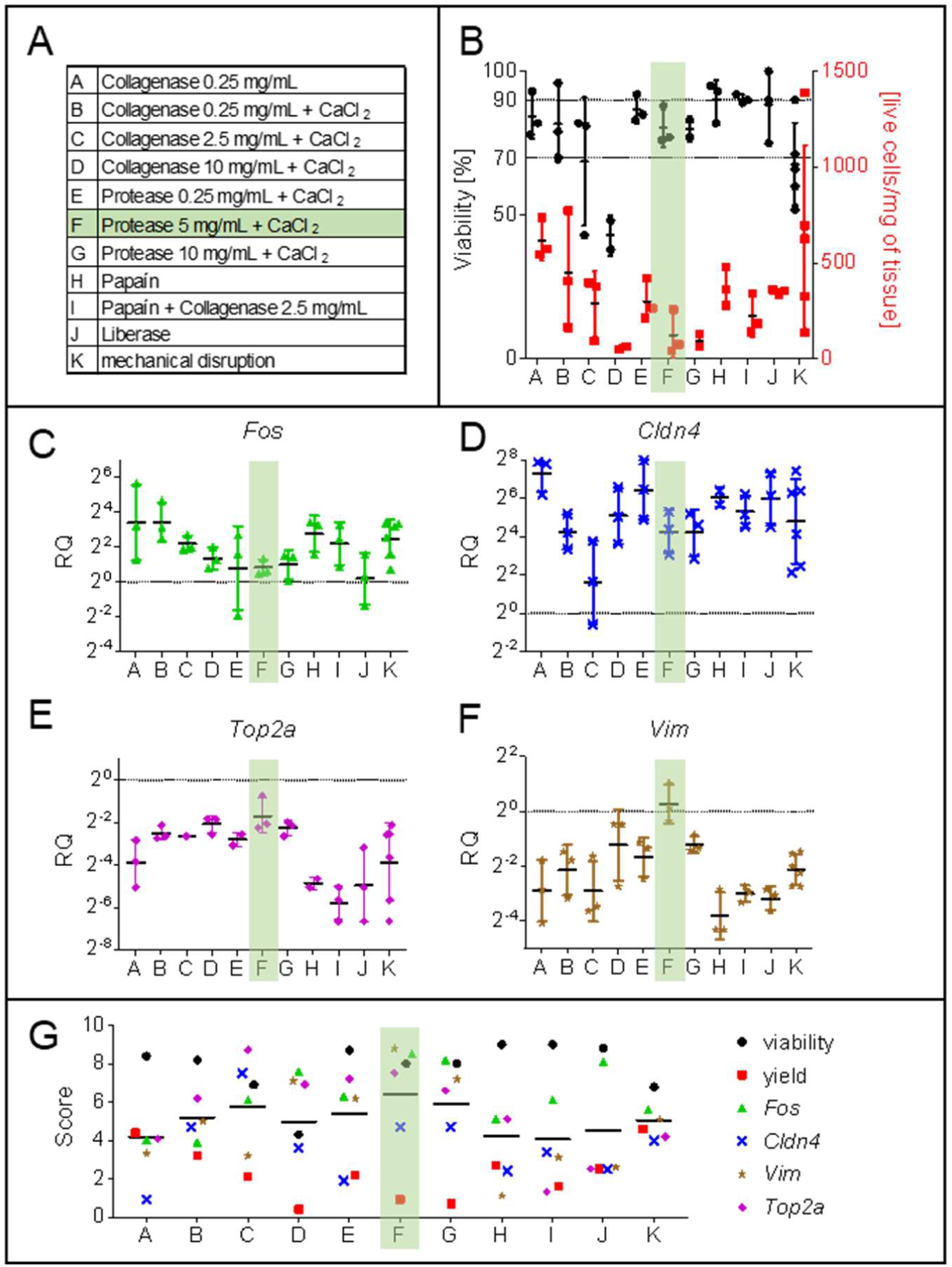
The effect of various enzymes and their combination on dissociation of the tumour tissue. (**A**) List of compared conditions. The condition found as optimal is marked in green. (**B**) Effect of using different enzymes during tumour dissociation (4 °C for 1 hour) on cellular viability (black) and yield (red). The recommended viability at least 90 % and minimal viability 70 % are marked in the graph. (mean ± SD) (**C-F**) The effect of different enzymes and their concentrations on relative gene expression of marker genes after dissociation normalized to non-dissociated tissue piece. The black line represents the expression in non-dissociated tumour tissue. (geometric mean with geometric SD) (**C**) *Fos* – a member of IEGs, (**D**) *Cldn4* – a marker of tumour endothelial stem cells, (**E**) *Top2a* – a marker of proliferating cells, (**F**) *Vim* – a marker of tumour epithelial cells. (**G**) Score (from worst 0 to best 10) were calculated for each condition and parameter. Mean of the scores is shown. See also **Figure S1**.

The first tested parameter was cell viability. A high cell viability is a critical parameter for all droplet based scRNA-Seq methods, because dying cells release RNA into solution and this can result in unspecific RNA background in all cells and decrease the sensitivity to identify different cell types. The manufacturers of single-cell droplet instruments have declared that an optimal cell viability should be more than 90-95 % (Illumina and Bio-Rad, 2017). Some manuals have suggested as low as at least 70 % viable cells (10x Genomics, 2018). In our experiment, none of the dissociation conditions resulted in cell viability higher than 90 % (**Figure 2B**). However, only two conditions showed an average viability lower than 70 %: Collagenase at the highest concentration (10 mg/ml) and mechanical disruption.

The second studied parameter was the cell yield after dissociation. The amount of the tissue used can also be a limiting parameter, especially for smaller and younger tumours. The low number of live cells obtained during dissociation indicates an ineffective dissociation or a high proportion of the disrupted cells. Surprisingly, the highest yield was achieved after mechanical disruption and dissociation using the Collagenase (0.25 mg/ml) without the addition of CaCl_2_ (**Figure 2B**). The higher enzymatic activity (after addition of CaCl_2_ or at higher concentrations of the enzymes) probably led to the disruption of cell membranes, which resulted in lower dissociation yields.

Any changes of the gene expression during sample preparation are unwanted. The artificial induction of gene expression and specific cell type dissociation were measured using RT-qPCR analysis of IEGs and mammary tumour cell types markers (**Figure 2C-F, Figure S1**). Gene expression was compared between the single-cell suspension and the non-dissociated tissue sample prepared in parallel (**Figure 1**). Activation of IEGs was determined using expression of *Fos*, as a prime example of IEG (**Figure 2C**). Expression most closest to the non-dissociated tissue was obtained by using Liberase, Protease (any concentration) and Collagenase (10 mg/ml). The remaining conditions led to an increase in *Fos* expression more than 5 fold relative to the non-dissociated tumour. The marker of cells, which easily dissociate from tumour tissue into suspension (*Cldn4*+; tumour endothelial stem cells) was enriched in all studied conditions (**Figure 2D**). On the other hand, the expression of a marker of cells which hardly dissociates from tumour tissue into suspension (*Vim*; mammary tumour epithelial cells) or a marker of proliferating cells (*Top2a*) was decreased (**Figure 2E,F**). The best condition that exhibited comparable expression relative to the non-dissociated tissue were observed using Protease and Collagenase with the addition of CaCl_2_ (**Figure 2E**). Similarly, the smallest relative expressional changes of *Vim* (**Figure 2F**) were determined in Collagenase (10 mg/ml) and Protease treatments (any concentration).

A scoring system from 0 (worst) to 10 (best) was developed and applied to combine all studied parameters (details in Method section). The lowest score (6.4 ± 3.1), which reflected the most efficient and reliable dissociation parameters, was obtained for Protease from *Bacillus licheniformis* at a concentration of 5 mg/ml (**Figure 2G**). The remaining conditions showed scores ranging from 4.1 to 5.9.

### The effect of the centrifugal force on the quality of cell suspension

The decrease in the gene expression of the cellular markers such as *Vim* and *Top2a* (**Figure 2C-F**), suggested a loss of these particular cell subpopulations in the final suspension. Previous studies have recommended the use of a higher centrifugal force for the collection of smaller cells. We tested centrifugation forces ranging from 300 to 3000 g (**Figure 1**) in combination with Protease (5 mg/ml) for dissociation (4 °C, ActD) and analysed the same set of parameters as in **Figure 2**. We found that cell viability is negatively affected by an increased centrifugal force. (**Figure 3B**). On the other hand, the yield of cell collection is proportional to centrifugal force (**Figure 3C**). No effect on the expression of *Fos* was observed when the centrifugal force was increased (**Figure 3D**). Similarly, no changes were detected in the expression of the proliferation marker (*Top2a*) or mammary tumour epithelial cell marker (*Vim*) (**Figure 3F**,**G**). The positive effect of an increased centrifugation force was however observed on the expression of *Cldn4* (marker of tumour endothelial stem cells). The higher centrifugal force led to lower enrichment of the *Cldn4*+ cells (**Figure 3E**).

**Figure 3:**
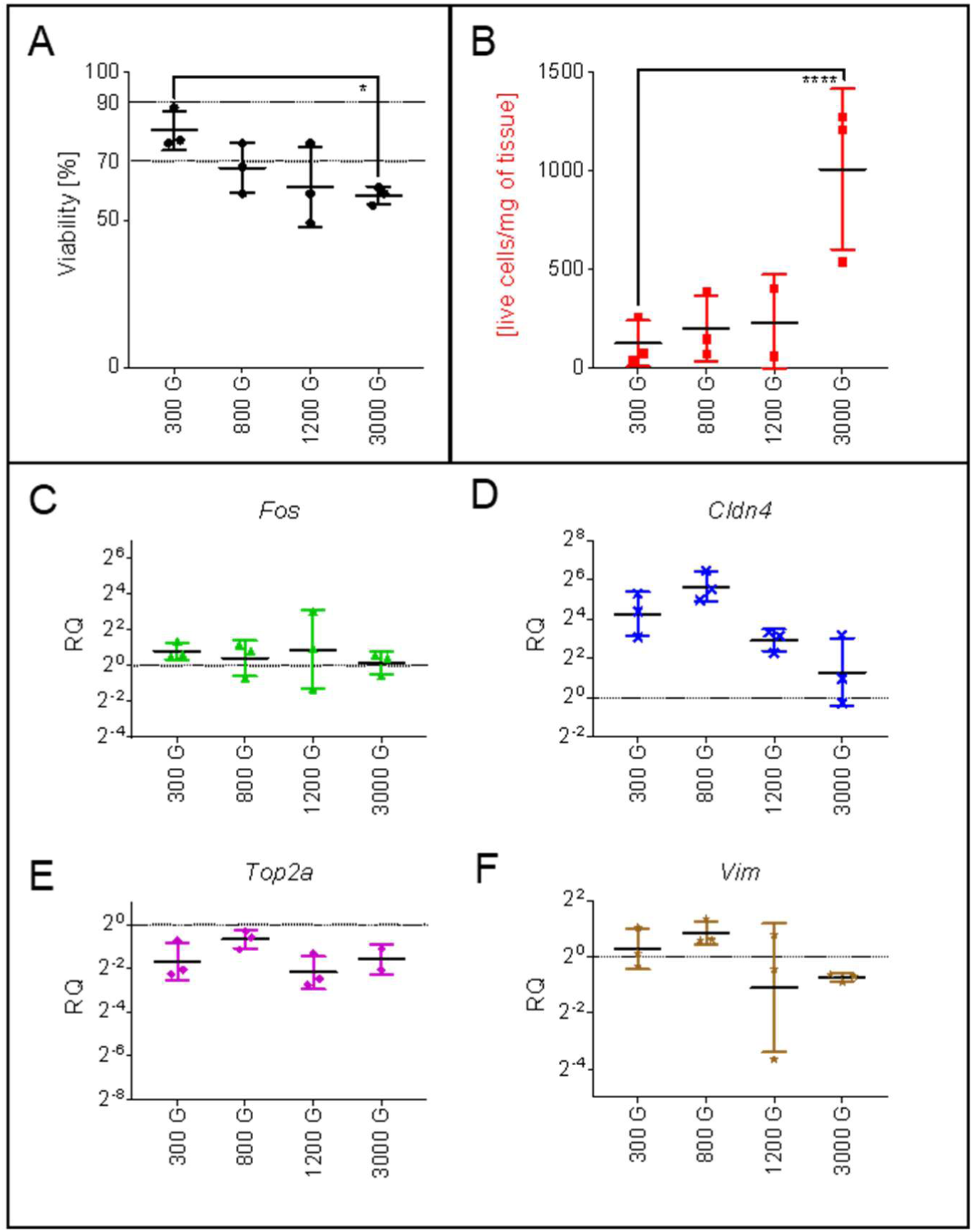
The effect of centrifugation force after tumour tissue dissociation on cellular viability and gene expression. (**A**) Cell viability after spinning them by different centrifugation force are shown in black. The recommended viability at least 90 % and minimal viability 70 % are marked in the graph. (mean ± SD) (**B**) Yields of cells after usage of various centrifugation forces are shown in red (mean ± SD). (**C-F**) The effect of centrifugation forces studied by relative expression of marker genes after dissociation normalized to non-dissociated tissue pieces. The black line represents the expression in non-dissociated tumour tissue. (geometric mean with geometric SD) (**C**) *Fos* - a member of IEGs, (**D**) *Cldn4* – a marker of tumour endothelial stem cells, (**E**) *Top2a* –a marker of proliferating cells, (**F**) *Vim* – a marker of tumour epithelial cells. Dunnett’s multiple comparisons test, **** - p <.0001, * - p < .05, n.s. - p > .05 See also **Figure S2**.

Even though the highest centrifugal force of 3000 g led to the highest yields and showed the best preservation of the cell composition, viability as a crucial parameter in single cell analysis was compromised. We found that centrifugation at 800 g resulted in an average viability of 70 %, which is the minimal required viability for further downstream analyses, while retaining appreciable cell yield and good composition of cell types.

### The effect of different temperature on the cell quality and expression of IEGs

Previous studies (Adam, Potter and Potter, 2017; Wu et al., 2017) have focused only on the impact of the activation of IEGs expression during tissue dissociation. We compared the effect of different dissociation temperatures (37 °C and 4 °C) and the presence of the transcription inhibitor (ActD) on the studied parameters (viability, yield, changes in cell types proportion), in combination with Protease (5 mg/ml) for dissociation and 800 g for centrifugation. Conditions were compared to dissociation at 37 °C in the absence of the inhibitor, because standard working temperature for the majority of the enzymes used for tissue dissociation is usually at 37 °C or higher.

Cell viability in samples dissociated at 37 °C in the absence of the inhibitor was 86 ± 5 % (**Figure 4A**). Although the treatment with ActD as well as the treatment at the lower dissociation temperature led to lower viability, the change was statistically insignificant relative to the control. However, we expected that the administration of ActD would lead to an increase in the number of dying cells (its toxic effect published by Cortes *et al*. (2016)), and subsequently to lower cell yield. Surprisingly, the yield was comparable or even higher during dissociation at 37 °C (**Figure 4B**). To explain this effect, we analysed changes in the number of the cells during incubation of the cell suspension at 37 °C in the presence or the absence of the ActD (**Figure S3A**). Incubation of the cells in the absence of the inhibitor led to a decreased cell number, but addition of the inhibitor stopped cells dying. The dissociation at 4 °C led to a decrease in the yield of the cells (**Figure 4B**). The observed difference in the yield between dissociation temperatures was greater after dissociation of the early stage tumour (10 days after injection) (**Figure S3B**).

**Figure 4:**
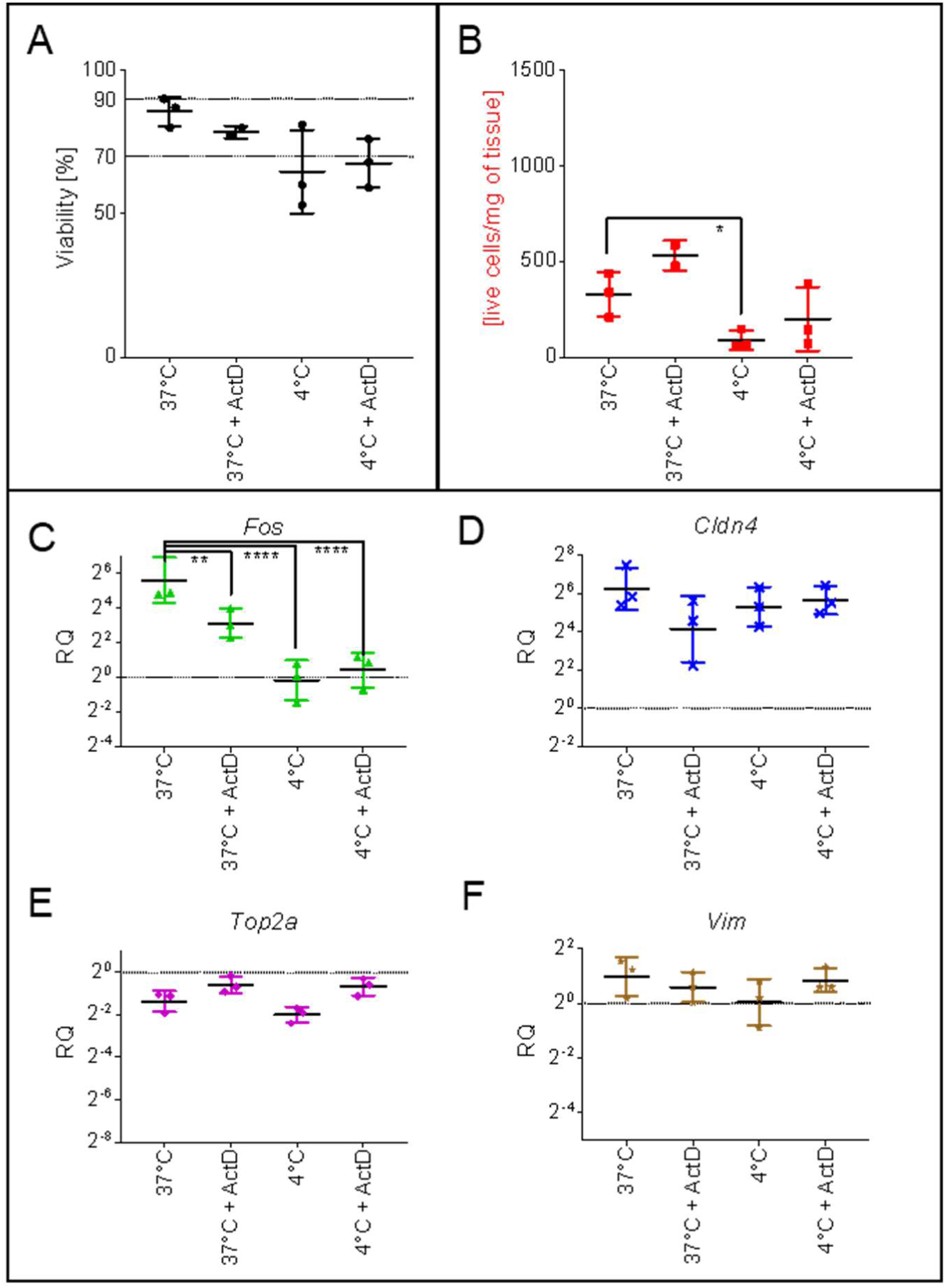
The effect of temperature oncell viability, yield and gene expression in dissociated tumour tissue. (**A**) Cell viability at different temperature and in the presence or absence of ActD are shown in black. The recommended viability at least 90 % and minimal viability 70 % are marked in the graph. (mean ± SD) (**B**) Yields of cells at different temperature and in the presence or absence of ActD are shown in red (mean ± SD). (**C-F**) The effect of dissociation temperatures and presence of ActD studied by relative expression of marker genes after dissociation normalized to non-dissociated tissue piece. The black line represents the expression in non-dissociated tumour tissue. (geometric mean with geometric SD) (**C**) *Fos* – a member of IEGs, (**D**) *Cldn4* – a marker of tumour endothelial stem cells, (**E**) *Top2a* – a marker of proliferating cells, (**F**) *Vim* – a marker of tumour epithelial cells. Dunnett’s multiple comparisons test, **** - p <.0001, ** - p < .01, * - p < .05, n.s. - p > .05 See also **Figure S3**.

Even though the dissociation at 4 °C resulted in a lower yield and lower cell viability, it more importantly resulted in an overall lower artificial induction of the IEGs. The fold change of *Fos* expression during dissociation at 37 °C was 48 *÷ 2.5 times higher (**Figure 4C**) in comparison with non-dissociated tumour tissue and 8.5 *÷ 1.8 times higher when ActD was used. The dissociation at 4 °C showed almost no change of *Fos* expression during dissociation, and the additional effect of ActD was minimal. No statistically significant changes of other gene expression markers were observed (**Figure 4D-F, Figure S3C-F**).

### The effect of the different dissociation conditions on the interpretation of scRNA-Seq experiment

We performed single-cell RNA-Seq to study the effect of temperature and the transcriptional inhibitor ActD, on the representation of individual cellular populations within the tumour sample (**Figure 5A**). We collected tumour tissue after 10 days of the growth, dissociated it and compared scRNA-Seq data with analysis using RT-qPCR from the same samples (**Figure 5A**). ScRNA-Seq analysis contained 654 cells, which were divided into four main cellular types: Tumour cells, Macrophages, T cells and Natural killer (NK) cells (**Figure 5B**).

**Figure 5:**
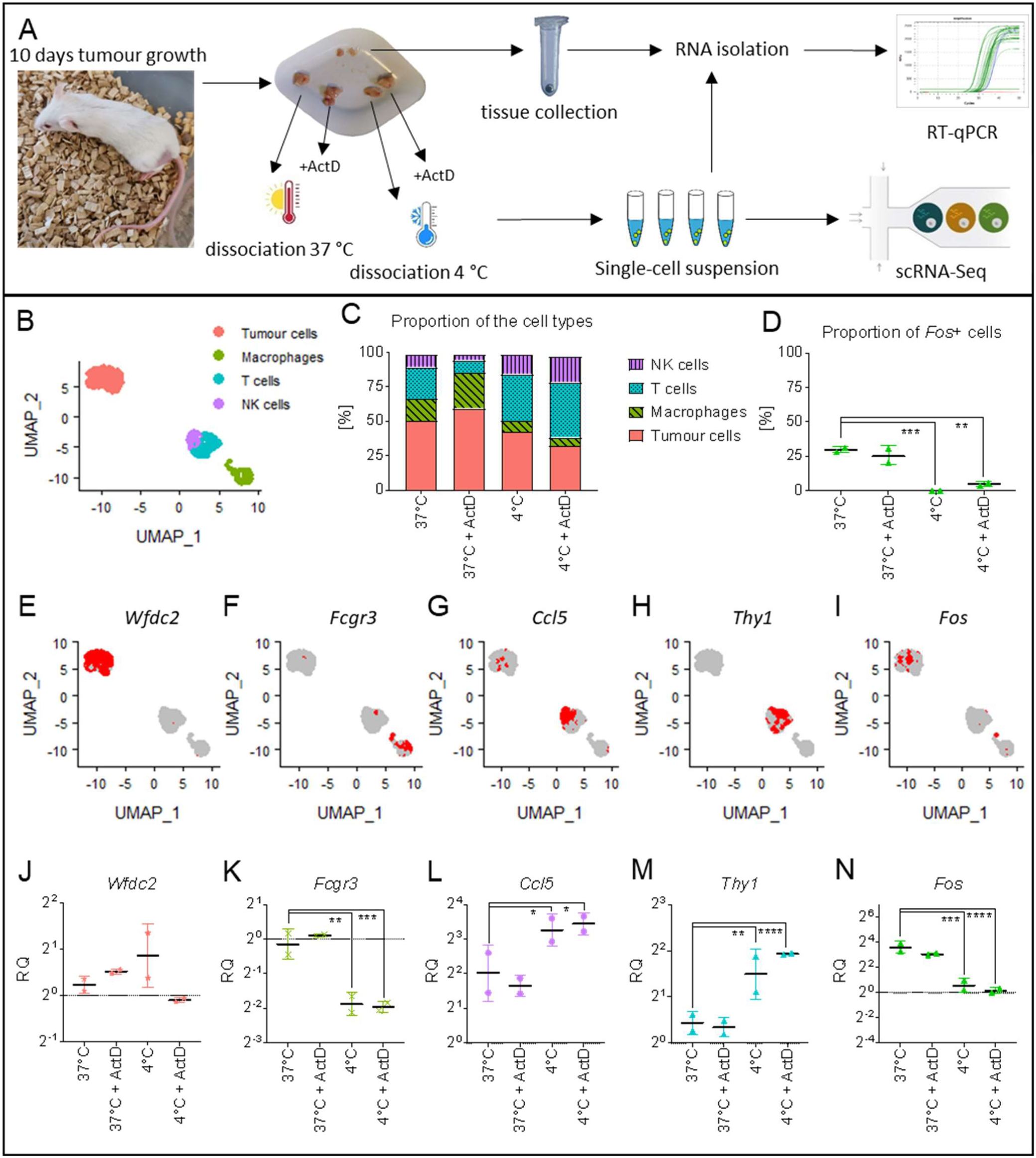
Comparison of scRNA and RT-qPCR for identification of population distribution in tumour tissue. (**A**) Scheme of the comparison of scRNA-Seq and RT-qPCR experiments. (**B**-**I**) Results from scRNA-Seq. (**B**) UMAP plot of identified cell types. (**C**) Proportion of the cell types after each different dissociation condition relative to all cells. (**D**) Proportion of the *Fos+* cells. (**E**-**I**) UMAP plot with marked individual cells positive for expression of different markers. (**J**-**N**) RT-qPCR analysis of relative expression of marker genes for cell population after dissociation of tumour compared with non-dissociated tissue. The black line represents the expression in non-dissociated tumour tissue. (geometric mean with geometric SD) (**D**,**I**) *Wfdc2* –a tumour cells (**E**,**J**) *Fcgr3* - Macrophages/monocytes (**F**,**K**) *Ccl5* - NK cells (**G**,**L**) *Thy1* - T cells (**H**,**M**) *Fos* – one member of IEGs. Dunnett’s multiple comparisons test. **** - p < .0001, *** - p < .001, ** - p < .01, * - p < .05, n.s. - p > .05

ScRNA-Seq analysis of the cell type representation showed a higher proportion of the Tumour cells and Macrophages after dissociation at 37 °C while a higher proportion of the NK and T cells after dissociation at 4 °C (**Figure 5C**). To compare the scRNA-Seq results with data from the original non-dissociated tissue, we analysed the expression of cell markers of these populations using RT-qPCR. The distribution of the cell populations showed minimal or small differences in the expression of all four markers after dissociation at 37 °C, but higher differences in the expression of markers of the Macrophages, NK cells and T cells after dissociation at 4 °C (**Figure 5J-M**). The effect of the ActD on the population representation was minimal. The induction of IEGs expression was analysed as changes of the expression of *Fos*. The analysis revealed that 30 % of examined cells were positive for the artificial induction of *Fos* gene expression when dissociated at 37 °C compared to only 5 % when dissociated at 4 °C. (**Figure 5D**). Similar results were observed after RT-qPCR (**Figure 5N**). Interestingly, the activation of *Fos* expression was specific mainly to tumour cells (**Figure S4A,C**) as 53 % of tumour cells showed expression of *Fos* at 37 °C dissociation (**Figure S4B**). Even though the dissociation at 37 °C led to better preservation of the cell composition, the crucial reduction of artificial gene expression remains the main benefit of lower dissociation temperature.

## DISCUSSION

The ability to dissociate tissue into a suspension of individual cells, while preserving the cell composition and initial gene expression as the original tissue, is required for precise single cell analyses. Maintaining such consistency becomes exceptionally vital for the accurate description of the tissue microenvironment and heterogeneity as well as the identification of new cell types and the characterization of the cellular behaviour within complex biological tissues. The protocol for tissue dissociation is complex and each step positively or negatively affects the quality and interpretation of the experiment. In our study, we compared several factors such as the selection of the dissociation enzyme, dissociation temperature, presence of transcription inhibitor and centrifugal force on the tumour tissue dissociation and single-cell suspension quality.

In our experiment, we tested several enzymes and their concentrations, with the goal to identify the optimum dissociation parameters to transform mouse tumour tissue into single cell suspension while maintaining its quality, for use with droplet based single cell library preparation instrument such as Chromium (10x Genomics) or ddSEQ (Bio-Rad). We created a scoring system to compare not only the dissociation yield and viability, but also to test the expressional changes of specific cell type markers and IEGs. Using our scoring system, we found that overall the best choice for 4T1 tumour dissociation was the use of the Protease from *Bacillus licheniformis*. Previously this protease has been deemed as “Psychrophilic” or “cold active” (Adam, Potter and Potter, 2017; Joshi and Satyanarayana, 2013), even though Tokoyawa *et al*. (2010) described its maximal activity at 50 °C. A decrease in temperature led to a decrease in the protease activity: 20 % of maximal activity at 37 °C, and only about 5 % of maximal activity at 20 °C (Toyokawa et al., 2010). The problem with low activity of the enzyme at the low temperature was solved by using a higher concentration, which resulted in a better score in more cases. Comparing the relation between the expressional changes and the concentration of enzymes, we concluded that at low temperature, effective tissue dissociation is less dependent on the enzyme type but rather on the concentration used.

During tissue dissociation obtaining both a high cell yield and cell viability are important, but often contradictory factors. Lower dissociation temperature is required for inactivation of IEGs, but leads to the reduction of the enzymatic activity. In contrast, too high enzyme concentration can lead to cell membrane degradation, decreasing cell viability. Cell disruption rate is probably cell type dependent. Therefore, it is possible that an uneven degradation can result in a decrease or even loss of specific cell subpopulations. The optimal combination of parameters depends on the particular experimental requirements (cell concentration, minimal required cell viability and quality of cell suspension) and on the type and size of the dissociated tissue. For example, the low dissociation temperature led to changes in the proportion of the immune cells in the tumour microenvironment (**Figure 5C,J-L**). Such proportional changes caused by the dissociation protocol could led to an erroneous biological interpretations. This can be particularly problematic when monitoring the immune response during cancer progression or during immunotherapy (Miragaia et al., 2019).

On the other hand, the higher temperature during dissociation led to the activation in the expression of IEGs, which can also result in the misinterpretation of the results. Members of IEGs are often used as cancer related markers, for example the *Fos* gene is considered as a prognosis marker for tumour progression in ovarian carcinoma (Mahner et al., 2008). Cell populations within the dissociated tumour samples have also been previously identified based on their IEGs expression when using scRNA-Seq experiments (Freeman, Jung and Ogle, 2016; Puram et al., 2017; Crinier, A. et al., 2018). However, the majority of these studies provided no thorough verification of their results using an alternative method, and therefore it is possible that their conclusions may be affected by artificial induction of IEGs expression. We found only one similar study by Tirosh *et al*. (2016), which confirmed scRNA-Seq results using immunohistochemistry.

We tested the effect of different factors on IEGs and found that the main source of induced expression is the dissociation temperature (**Figure 2-4**). In our hands, the addition of ActD reduced IEGs activation. However the improvement compared to the low dissociation temperature was negligible (**Figure 4**) and only the combination of both approaches led to the complete inhibition of artificial IEGs expression. Another source of cell stress comes from centrifugation and we hypothesized that the higher centrifgual force will lead to a higher effect on the IEGs expression. In contrast to our expectations, we observed no changes in the IEGs expression dependent on the centrifugal force (**Figure 3**). We believe that additional factors, which include centrifugation at 4 °C and using a minimal acceleration and breaking of the centrifugal speed, reduced the stress response.

The centrifugal force is an important factor for cell viability too. Optimization is required especially when working with fragile and sensitive cells. On the other hand, smaller/lighter cells require higher centrifugal forces for their efficient capture (Moroz, 1984). We performed a literature search and found only one example where the optimization of centrifugal force was discussed (10x Genomics, 2017). Even though the effect of the centrifugal force during single-cell suspension preparation is considerable for its quality, it is usually performed without optimization and often the published information about centrifugation is missing in the Methods section (Patel et al., 2014; Chung et al., 2017; Winterhoff et al., 2017). Standard protocol recommends a centrifugal force of 300 g (Miltenyi Biotec, 2018), even though a recently published protocol used a centrifugal force up to 1500 g (Bykov, Kim and Zamarin, 2019). Nevertheless, the choice for the optimal force is relies on the compromise between viability, yield and the potential capturing of all the cell types.

We summarized the effects of the different factors on the quality of single-cell suspension in **Table 1** and discussed the other potential factors and their expected impact in **Table 2**. We showed that optimization of the dissociation protocol is a complicated and laborious process and that there is no gold standard applicable for every tissue type and biological experiment. Improvement of single or multiple parameters can result in changes in other parameters and subsequently on the quality of single-cell suspension. Surprisingly using the same condition showed good reproducibility even though the samples were collected from tumour tissue, which is known for its large heterogeneity.

**Table 1:** Effects of the different tested factors used during single-cell suspension preparation that affects the quality of the single-cell suspension.

|  | Higher enzyme concentration | Higher temperature | Presence of the inhibitor of gene expression | Higher centrifugation force |
| --- | --- | --- | --- | --- |
| Positive effect | Higher dissociation effectivity | Higher dissociation effectivity | Inhibition of the changes of the cell state.<br>Higher yield at 37 °C | Better capture of all the cell types<br>Higher yield |
| Negative effect | Lower yield | Activation of IEGs | Not observed | Lower viability<br>Disruption of fragile cells |

**Table 2:**
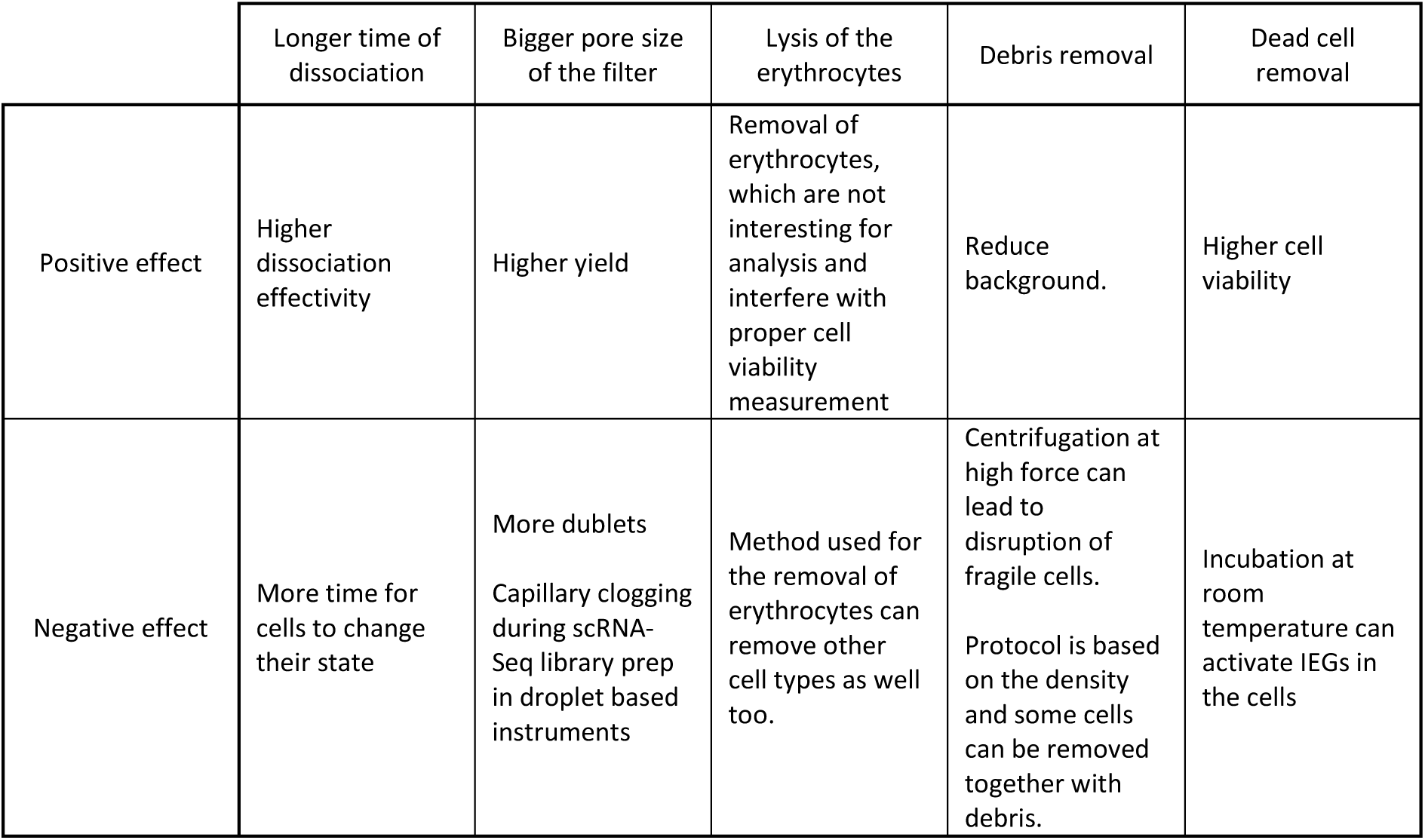
Potential effects of the other factors used during single-cell suspension preparation that may affect the quality of the single-cell suspension.

## METHODS

### Ethics statement

All animal studies were approved by the Czech Academy of Sciences and conducted in accordance with the Czech Council guidelines for the Care and Use of Animals in Research and Teaching.

### Tissue culture and mice

The mouse 4T1 breast cancer cell line was obtained from the American Type Culture Collection (ATCC CRL-2539) and was routinely cultivated in Roswell Park Memorial Institute (RPMI, Sigma) medium supplemented with 10 % fetal bovine serum (Thermo Scientific) and 100 U/mL penicillin, 100 µg/mL streptomycin; in 5 % CO_2_ and 37°C. When injected into Balb/c mice, 4T1 spontaneously produces highly metastatic tumour. Balb/c mice were subcutaneously injected with 1×10^6^ of 4T1 cells. After 20-25 days, mice were sacrificed, and tumours were collected for further single cells preparation. Biological triplicate were used for every experiment.

### Single-cell suspension preparation

Each tumour was cut into several pieces. One piece from each tumour was used for control RNA isolation of non-dissociated tissue. Remaining pieces of tumours were weighted (about 100 mg), minced by using polypropylene Pellet pestles for 1.5 mL tubes (Sigma, Z359947) in Eppendorf tubes and transferred into gentleMACS C tubes (Miltenyi Biotec) containing dissociation solution composed of 5 mL of RPMI media with tested enzymes according to **Table 3**. To inhibit the transcription during dissociation, we used 2 nM (25 μg/ml) of Actinomycin D (ActD, Sigma A1410) in dissociation solution and 0.2 nM ActD (2,5 μg/ml) in all following solutions. Tubes were mixed by using gentleMACS Dissociator (Miltenyi Biotec) program m_impTumor_02.01 (about 30 s) placed in 4 °C and then on a rotating platform set at 20 rpm for 20 minutes in 4 °C. This step was repeated 3 times for a total of 1 hour of dissociation. Dissociation at 37 °C were performed using the gentleMACS OctoDissociator (Miltenyi Biotec) with heaters following the program parameters as outlined in **Table 4**. The total dissociation time was 1 hour.

**Table 3.** list of tested enzyme conditions

|  | Enzyme | Enzyme concentration | Medium | CaCl<sub>2</sub> | DNase |
| --- | --- | --- | --- | --- | --- |
| A | Collagenase | 0.25 mg/ml | RPMI | x | 50 µg/ml |
| B | Collagenase | 0.25 mg/ml | RPMI | 10 mM | 50 µg/ml |
| C | Collagenase | 2.5 mg/ml | RPMI | 10 mM | 50 µg/ml |
| D | Collagenase | 10 mg/ml | RPMI | 10 mM | 50 µg/ml |
| E | Protease | 0.25 mg/ml | RPMI | 10 mM | 50 µg/ml |
| F | Protease | 5 mg/ml | RPMI | 10 mM | 50 µg/ml |
| G | Protease | 10 mg/ml | RPMI | 10 mM | 50 µg/ml |
| H | Papain | 61.25 mg/ml | PBS -/-, EDTA (0,5 mM), L-Cystein (1 mM) | x | x |
| I | Papain<br>Collagenase | 61.25 mg/ml<br>2.5 mg/ml | PBS -/-, EDTA (0,5 mM), L-Cystein (1 mM) | x | 50 µg/ml |
| J | Liberase | 0.25 mg/ml | RPMI | x | 50 µg/ml |
| K | x | x | RPMI | x | 50 µg/ml |

Collagenase – Collagenase from *Clostridium histolyticum*, Sigma C7657, 250 mg/ml stock solution in PBS
Protease – Protease from *Bacillus licheniformis*, Sigma P5380, 250 mg/ml stock solution in PBS -/-
Liberase - Liberase™ TL Research Grade, Roche 5401020001, 2.5 mg/ml stock solution
Papain – Papain solution, Merck L2223
DNase – DNase I, Roche 11284932001, 10 mg/ml in nuclease free water (NFW, Invitrogen 10977-035)
CaCl<sub>2</sub> –1 M stock solution in NFW

**Table 4.** gentleMACS Octo Dissociator program for dissociation at 37°C

|  |
| --- |
| 1. ramp 200 rpm, 10" |
| 2. loop 3x |
| 3. spin -200 rpm, 1" |
| 4. spin 200 rpm, 10" |
| 5. end loop |
| 6. temp ON |
| 7. loop 2x |
| 8. spin -30 rpm, 30' |
| 9. spin 200 rpm, 10" |
| 10. end loop |
| 11. temp OFF |
| 12. end |

Immediately after dissociation, all tubes were kept on ice while all solutions were pre-chilled in the ice. Cell suspension was filtered through a 30 µm strainer (CellTrics, 04-004-2326) and from this point kept on ice. Cells were spun down (5 min, 800 g•, 2 °C, minimal acceleration and break, Eppendorf 5810 R with swing-×bucket rotor A-4-44). The medium was discarded and the cells were dissociated in 15 mL of ice cold ACK solution (0.15 M NH_4_Cl; 10 mM KHCO_3_; 0.1 mM EDTA; pH 7.3) to remove erythrocytes. The cells suspension were immediately spun (5 min, 800 g•, 2 °C, minimal acceleration and break) and resuspended in 3.1 mL of ice-cold PBS without Ca^2+^ and Mg^2+^ (PBS -/-, Sigma, D8537). 900 µL of ice-cold debris removal solution (Miltenyi Biotec, 130-109-398) was added to the solution, mixed, carefully overlaid with 4 mL of PBS -/- and spun (10 min, 3000 g, 2 °C, full acceleration and brake). 5 mL of the upper phase solution was removed, the falcon tube was filled up with PBS -/-, mixed by inverting 3 times and spun (5 min, 800 g•, 2 °C, minimal acceleration and break). The supernatant was removed and dead cells were removed using the Dead Cells Removal Kit according to the manufacturer’s instruction (Miltenyi Biotec, 130-090-101). The collected effluent was spun (5 min, 800 g•, 2 °C, minimal acceleration and break) and the cells resuspended in 110 μl of PBS -/-. 10 μl of the cells suspension were mixed with 10 μl of Trypan Blue (Bio-Rad, #1450021) and the number of the cells and viability were analysed using BioRad TC20 cell counter. The remaining 100 μl of cells suspension were transferred to 1.5 ml tube, immediately frozen using dry ice and stored in the −80 °C freezer.

- For testing of effect of different centrifugation force, centrifugation force between 300 – 3000 g were used.

### Gene expression analysis

#### RNA-isolation

Tumour tissue were collected into 2 ml tubes with pre-chilled steel beads (Qiagen Stainless Steel Beads, 5 mm, 69989) and stored at −80 °C freezer. Cell suspension were collected into 1.5 ml tubes, frozen on dry ice and stored at −80 °C freezer. Samples were mixed in 1 ml of TRI Reagent (Sigma-Aldrich, T9424) with 2 μl of GlycoBlue (AM9515, Invitrogen). Tissue were homogenized immediately using TissueLyser LT (Qiagen), for 10 minutes at 50 Hz and cell suspension were vortexed for 3 minutes. Total RNA was isolated following the manufacturer’s manual. The RNA pellet was dissolved in 50 μl of 1xTE buffer (Invitrogen, 12090-015) and reprecipitated with addition of 50 μl of 8 M LiCl (Sigma-Aldrich, L7026) at −20 °C freezer overnight. This solution were centrifuged for 30 minutes at 16000 g. The supernatant were removed and RNA was washed twice with 1 ml of 80 % ethanol followed by centrifugation for 30 minutes. The final RNA pellet was diluted in 20 μl of 1xTE buffer (or 100 μl for samples from non-dissociated tissue). The concentration of RNA from tissue were measured using Nanodrop 2000 (Thermo Scientific).

#### RT-qPCR

The RNA from the tumour tissue were diluted to a final concentration of 5 ng/μl while the RNA from the cell suspension was used undiluted. The 5 μl of RNA were mixed with 0.5 μl RNA Spike I (TATAA, Biocenter), 2 μl of TATAA GrandScript cDNA Synthesis Kit and 2.5 μl of 1xTE Buffer. Reverse transcription was performed according to the manufacturer’s protocol (22 °C 5 min, 42 °C 30 min, 85 °C 5 min, 4 °C) and the synthesized cDNA was diluted 10 times in 1xTE buffer. 2 μl of the final diluted cDNA was added to the qPCR reaction (2x SYBRGreen mix, TATAA Biocenter, 400 nM primers mix and Nuclease-free water to final volume 7 µl). Primers sequences are listed in the **Table S1**. The qPCR were run on BioRad C1000 Thermal cycler and protocol for qPCR was: 1 minute at 95 °C; 50 cycles of 95 °C for 3 seconds, 60 °C for 30 seconds and 72 °C for 10 seconds; followed by melting curve analysis.

#### Data analysis and statistics

Measured C_q_ values for each gene were normalized to *Actb* (ΔC_q_). Relative quantity for each sample was calculated according to formula:

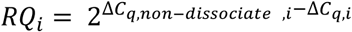

RQ_i_ – Relative quantity of expression of gene of replicate *i*

ΔC_q,non-dissociated,I_ - ΔC_q_ for non-dissociated control tissue sample of replicate *i*

ΔC_q,i_- ΔC_q_ for dissociated sample of replicate *i*

The final RQ is expressed as the geometric mean multiplied or divided by (*÷) geometric Standard deviation (geometric SD) (Kirkwood, 1979). The statistical significance was calculated from log2 transformed RQi values using GraphPad Prism 7 Two-way ANOVA with Dunnett’s multiple comparisons test.

The cell viability is expressed as the arithmetic mean plus minus (±) SD.

The yield of the live cells was calculated as number of live cells divided by weight of tumour tissue used for dissociation. The yield is expressed as arithmetic mean ± SD.

#### Scoring

Six different parameters were scored to choose the best condition:

1. Cell viability.
2. Number of live cells obtained from one milligram of the tissue.
3. Expression of four different genes (*Fos, Cldn4, Top2a, Vim*)

For each measured parameter were calculated score from 0 to 10 to choose the best condition, where 0 is the best and 10 is the worst.

##### For viability

0 is 0 % viability and 10 is 100 % viability

##### For number of live cells

0 is for 0 live cells and 10 is for 1500 or more live cells from mg of the tissue.

##### For gene expression

all numbers were log2 transformed and absolute values were calculated. Score 10 is for no changes between control tissue and dissociated sample (log_2_(RQ) = 0) and score 0 is for the highest change for tested gene.

### Single-cell RNA Sequencing

#### Cell suspension preparation

Balb/c mice were subcutaneously injected with 1×10^6^ of 4T1 cells. After 10 days, tumours were collected, weighted and processed according to the protocol mentioned above. A pool of 6 pieces of the tumour from different mice were used for each condition. In the final step, cells were resuspended in 100 μl PBS-/- with 0.04 % BSA (AM2616, Invitrogen) **without** ActD and concentration of the cells was analysed using TC20 cell counter (BioRad). The suspension was spun (5 min, 800 g, 2 °C, minimal acceleration and break), supernatant removed and cells were resuspended to a final concentration of 2500 cell/μl. The concentration and viability of the cells were measured two times and marked in the table (**Table S2**).

#### Library preparation

Two technical duplicates per condition were prepared (in total 8 samples). ScRNA-Seq libraries were prepared using SureCell WTA 3’ Library Prep Kit for the ddSEQ System (Illumina/Bio-Rad) according to manufacturer’s manual. Briefly, cells were individually partitioned into the droplets together with beads with barcodes and RT mix using ddSEQ Single-Cell Isolator (Bio-Rad). The cell lysis and RT took place in each droplet. Then, because cDNA:mRNA from each cells is barcoded, droplets could be disrupted and second strand were synthesized in bulk. After second strand synthesis, double stranded cDNA were analysed using Fragment Analyzer (AATI, HS NGS Fragment kit, DNF-474).

Double-stranded cDNA were tagmented using Nextera SureCell transposome. Final libraries were subsequently indexed during PCR amplification and purified. Before pooling of final libraries, the quality of the libraries were measured using a Fragment Analyzer (AATI, HS NGS Fragment kit, DNF-474). The pooled libraries were sequenced using NextSeq 500 instrument (Illumina) in HighOutput mode with setup – Read1 70 cycles, Index1 8 cycles, Read2 88 cycles. Read 1 contains 0-5 bp long Phase Block, three 6 bp long cell barcodes (BC1-3) split with two linkers 15 bp long, cell barcodes are followed with ACG sequence, 8 bp long unique molecular identifiers (UMI), GAC sequence and several Ts. Read 2 contained the insert sequence in the same strand as the mRNA.

#### Data analysis

On average 30 M reads per sample were obtained. Count table per cell were generated using umi-tools (v. 1.0.0) (Smith, Heger and Sudbery, 2017) following the manual with small modifications. At first, umi_tools whitelist was used to identify a list of correct cell barcodes with bc-pattern defined as regex (full command available in **File S1**) and set-cell-number to 3000. Next, umi_tools extract was used to copy cell barcode and umi sequence from read1 to the name of the read2. The whitelist generated from the previous step was used to filter reads with correct cell barcode with option for error correction of mismatches in it. Read 2 was then used alone for further downstream analysis. The low quality reads and adaptor sequences were removed using TrimmomaticSE (Bolger, Lohse and Usadel, 2014) with the parametres “ILLUMINACLIP:/mnt/d/Adapters/ddSeq_SureCell.fa:2:30:10 LEADING:3 TRAILING:3 SLIDINGWINDOW:4:15 MINLEN:36”. Then, reads were mapped to the mouse genome (GRCm38) using STAR (v. 2.7.1a) (Dobin et al., 2013) with the removal of reads which were mapped to more than one site. Reads were then assigned to the genes using featureCounts (v. 1.6.0) (Liao, Smyth and Shi, 2014) using the annotation *genecode*.*vM8*.*gtf* and parameter “-s 1” for definition of strand specificity of reads. Finally, the obtained BAM file was sorted and indexed using samtools (v. 1.7) (Li et al., 2009; Li, 2011) and the final table with information about counts of UMI per gene per cell were generated using umi_tools count with parameters “—per-gene and –wide-format-cell-counts”. The full list of commands used for analysis is available as **File S1**.

#### Clustering

The processing of data were made using R (v. 3.5.2). The raw count tables from all samples were first loaded into a Single Cell Experiment object. Genes that were not detected across all samples, were removed. R package DropletUtils (v 1.2.2) (Lun et al., 2019) was used to identify empty drops in the dataset, and only “cells” with more than 200, less than 5000 UMIs and FDR ≤ 0,05 were processed in the next step. The Scater package (v 1.10.1) (McCarthy et al., 2017) was used to calculate the Quality control matrix, and features which were not detected in at least two cells were removed from the dataset. The data was then changed into a Seurat object and normalized (Seurat package v 3.0.2) (Stuart et al., 2019; Butler et al., 2018). Cells from technical replicates were analysed as one condition. UMAP clustering (v 0.3.9) (Becht et al., 2018; McInnes et al., 2018) with PCA reduction were used to identify cell clusters. Two clusters, which were identified as cell debris were removed from analysis and the new subset of cells were reclustered again. In total 654 cells were analysed (average 164 cells per condition, min 88 cells, max 219 cells). The markers of cell clusters were identified using the function “FindConservedMarkers” and clusters of similar cell type were merged into one. The full list of commands used for analysis is available as **File S1**.

## ACKNOWLEDGMENTS

We thank Lukáš Valihrach for helpful discussion during discussion about experiments; Ravindra Naraine for grammar corrections. We thank to Genomics and Bioinformatics Core Facility at Institute of Molecular Genetics of the Czech Academy of Sciences (RVO: 68378050) for borrowed Bio-Rad ddSEQ Single-Cell Isolator instrument. This work was supported by the Grant Agency of the Czech Republic GA17-24441S; Institutional support (RVO: 86652036 and BIOCEV CZ.1.05/1.1.00/02.0109).

## AUTHOR CONTRIBUTIONS

P.A. performed RNA isolation, RT-qPCR experiments, single-cell library preparation and data analysis, wrote the manuscript and prepared all figures. P.A., S.L. performed tissue dissociation S.L., J.T. performed cell preparation and animal surgeries M.K., J.T., R.S. conceived and supervised the study. All authors reviewed the manuscript.

## DECLARATION OF INTERESETS

The authors declare no competing of interests.

## SUPPLEMENTAL MATERIALS

**Figure S1:**
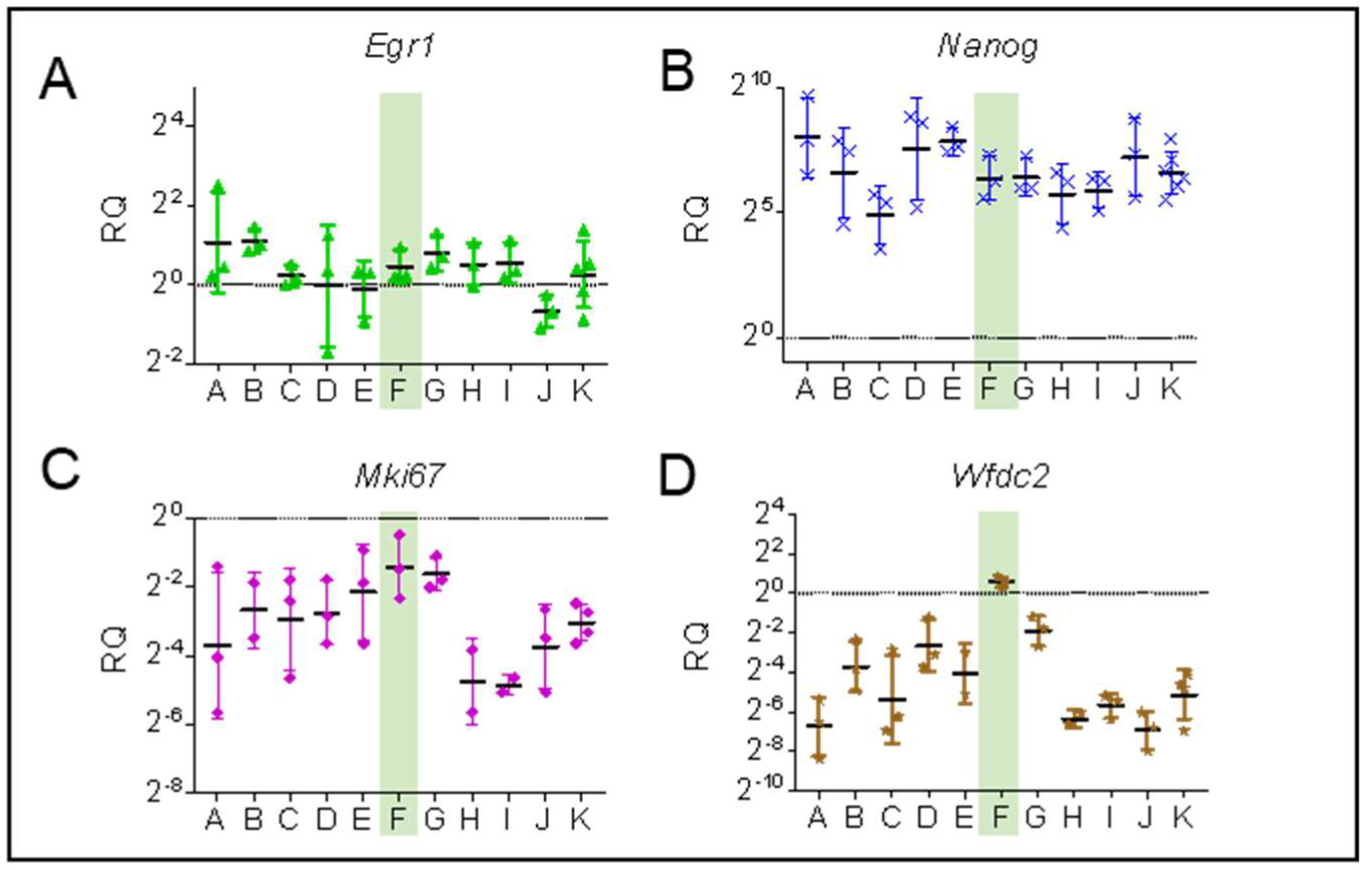
The effect of various enzymes and their combination on the dissociation of the tumour tissue. The effects of enzymes and their concentrations were assessed by comparing the relative expression of some additional marker genes after dissociation, normalized to the non-dissociated tissue pieces. The black line represents the expression in non-dissociated tumour tissue. (geometric mean with geometric SD) (**A**) *Egr1* – a member of IEGs, (**B**) *Nanog* – a marker of tumour endothelial stem cells, (**C**) *Mki67* – a marker of proliferating cells, (**D**) *Wfdc2* – a marker of tumour epithelial cells. The optimal dissociation condition marked in green.

**Figure S2:**
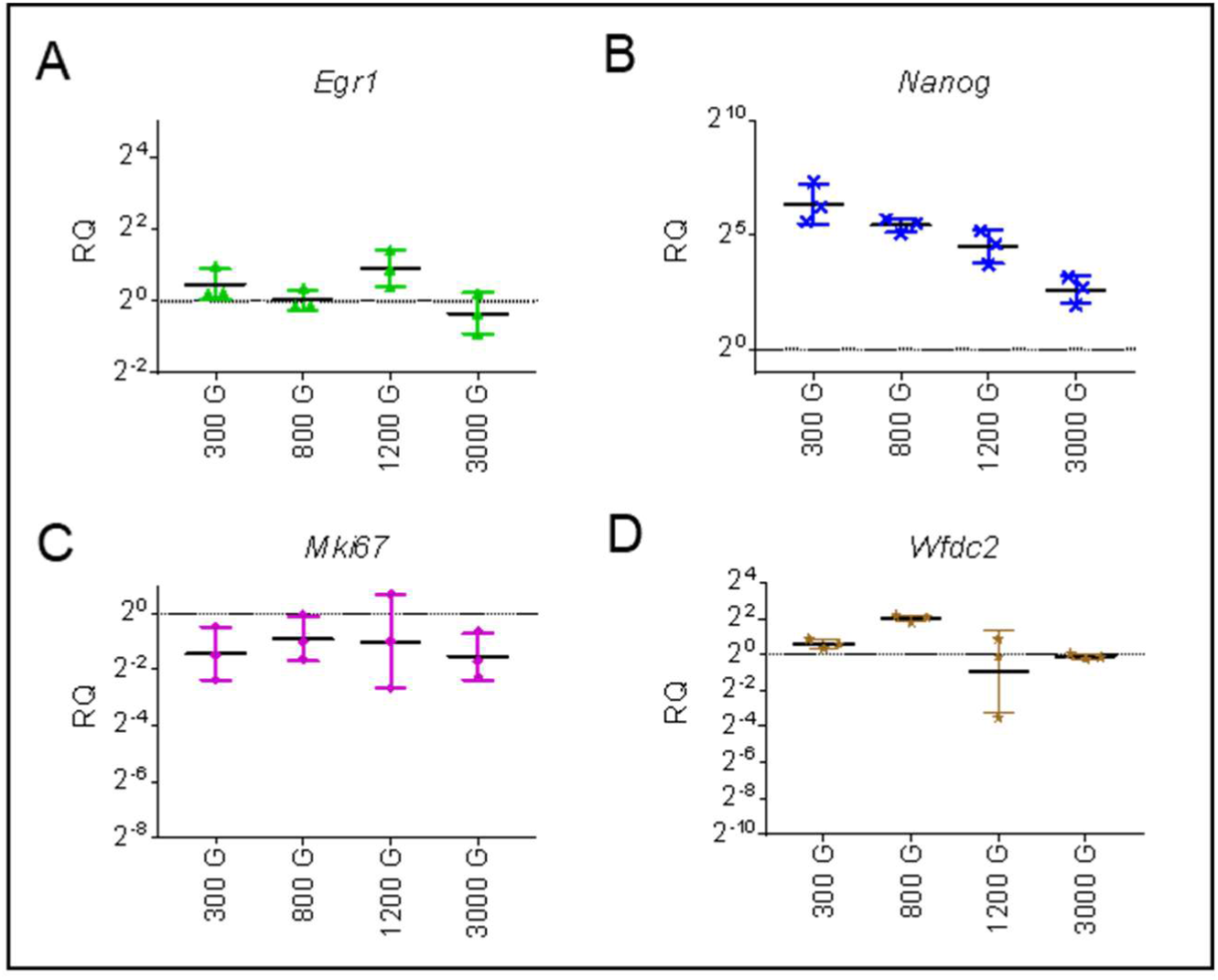
The effect of centrifugation force after tumour tissue dissociation on cellular viability and gene expression. Relative expression of additional marker genes after dissociation normalized to non-dissociated tissue piece comparing the effects of different centrifugation forces. The black line represents the expression in non-dissociated tumour tissue. (geometric mean with geometric SD) (**A**) *Egr1* – a member of IEGs, (**B**) *Nanog* – a marker of tumour endothelial stem cells, (**C**) *Mki67* – a marker of proliferating cells, (**D**) *Wfdc2* – a marker of tumour epithelial cells.

**Figure S3:**
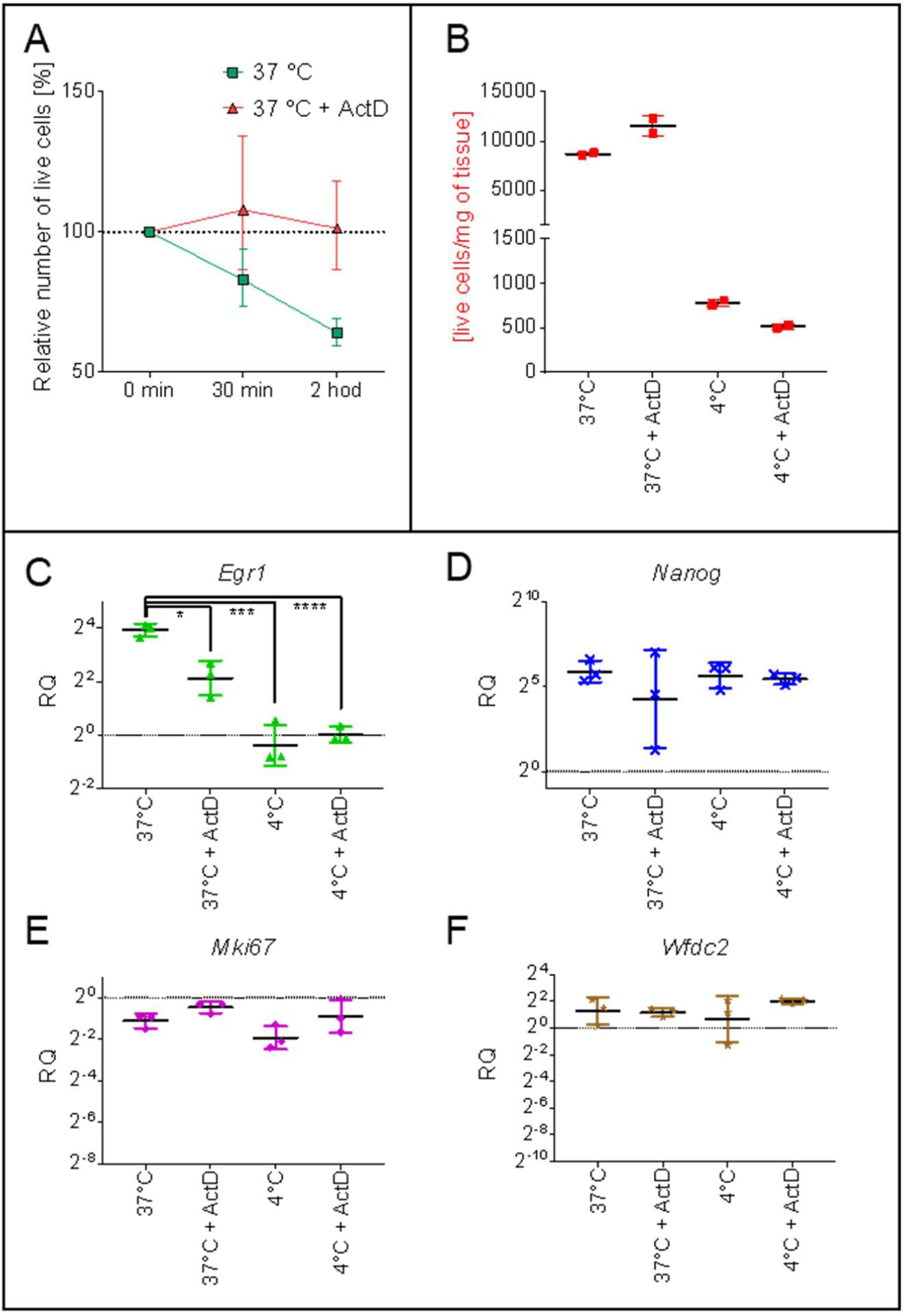
The effect of temperature and ActD on gene expression in dissociated tumour tissue. (**A**) Changes in number of live cells during incubation of cell suspension with or without ActD. Tumour tissue were dissociated at 37 °C for 1 hour with or without ActD. After that, cell suspension were washed and incubated at 37 °C in RPMI medium without enzyme with or without ActD. Number of the live cells were counted and compared with number of the live cells at the beginning of the incubation. (**B**) The yield of the live cells after dissociation of the tumour 10 days after injection. (**C-F**) Relative expression of additional marker genes after dissociation normalized to non-dissociated tissue piece comparing the effects of dissociation temperature and presence of ActD. The black line represents the expression in non-dissociated tumour tissue. (geometric mean with geometric SD) (**C**) *Egr1* – a member of IEGs, (**D**) *Nanog* – a marker of tumour endothelial stem cells, (**E**) *Mki67* – a marker of proliferating cells, (**F**) *Wfdc2* – a marker of tumour epithelial cells. The optimal dissociation condition marked in green. Dunnett’s multiple comparisons test, **** - p < 0,0001, ** - p < 0,01, * - p < 0,05, n.s. - p > 0,05

**Figure S4:**
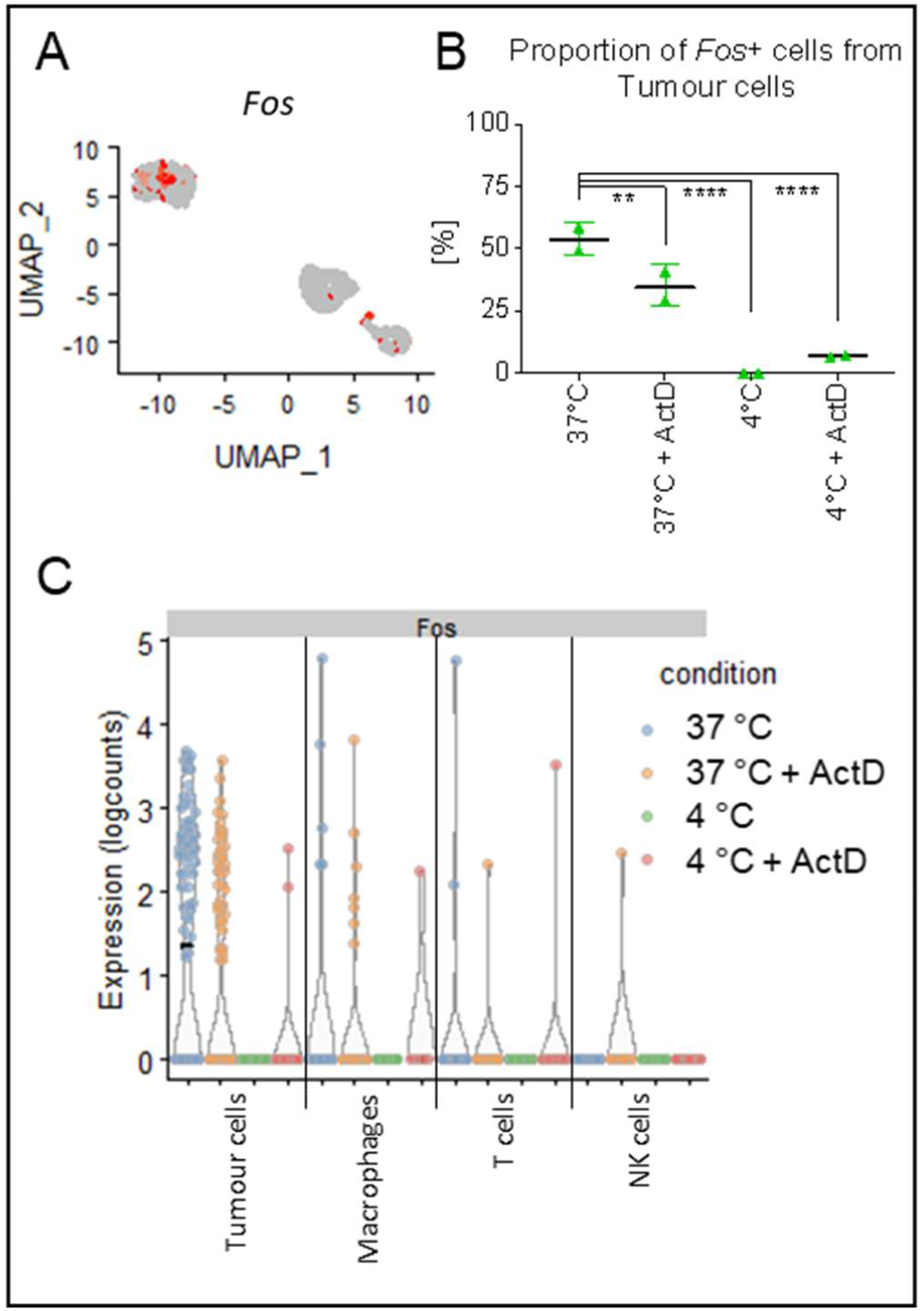
Identification of level of expression of *Fos* in tumour populations after different dissociation condition. (**A**) UMAP plot with marked individual cells positive for expression of *Fos*. (**B**) Proportion of *Fos+* cells from tumour cell type (mean ± SD). (**C**) Level of expression of *Fos* in individual cells of different cell types. Dunnett’s multiple comparisons test, **** - p < .0001, *** - p < .001, ** - p < .01

**Table S1:**
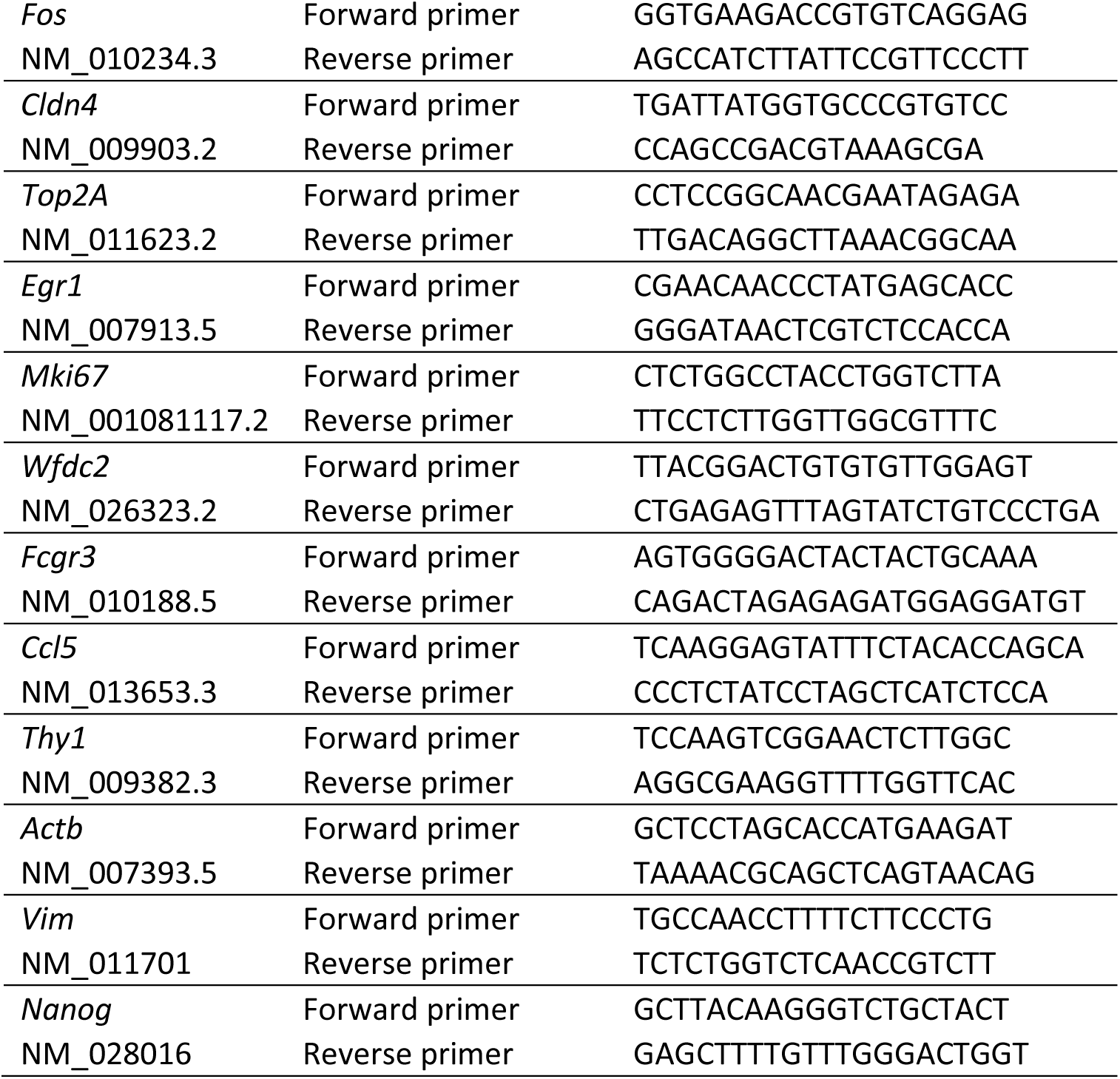
List of primers used for RT-qPCR

**Table S2:**
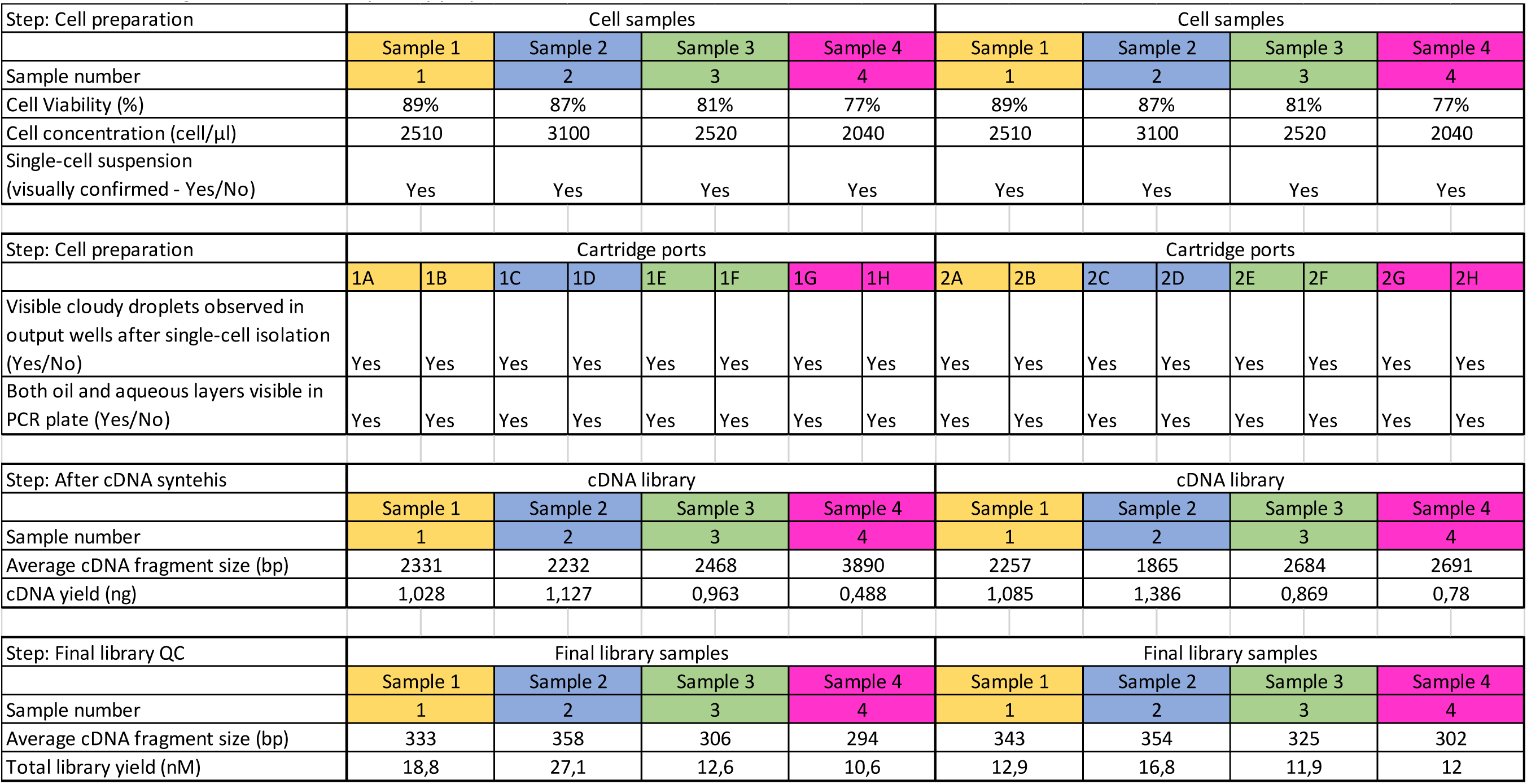
Lab tracking chart for scRNA-Seq library preparation

