## Supplemental file S1 for "Preparation of single-cell suspension from mouse breast cancer focusing on preservation of original cell state information and cell type composition"

### DATA PREPROCESSING

#### Creating of whitelist of cell barcodes

```
#UMI-tools version: 1.0.0
umi_tools whitelist \
--bc-pattern="(?P<discard_1>.{0,5})(?P<cell_1>.{6})(?P<discard_2>TAGCCATCGCATTGC){e<=1}
(?P<cell_2>.{6})(?P<discard_3>TACCTCTGAGCTGAA){e<=1}(?P<cell_3>.{6})(?P<discard_4>ACG)
(?P<umi_1>.{8})(?P<discard_5>GAC).*" \
--extract-method=regex --stdin raw/1A_1.fastq.gz \
--plot-prefix=whitelist/1A_whitelist --set-cell-number=20000 \
--log2stderr > whitelist/1A_whitelist.txt 2> log/1A_umi_whitelist.log;
```

#### Extraction of reads from the cells based on the whitelist and copying of the umi sequence and the cell barcode from read 1 into name of read 2

```
#UMI-tools version: 1.0.0
umi_tools extract \
--bc-pattern="(?P<discard_1>.{0,5})(?P<cell_1>.{6})(?P<discard_2>TAGCCATCGCATTGC){e<=1}
(?P<cell_2>.{6})(?P<discard_3>TACCTCTGAGCTGAA){e<=1}(?P<cell_3>.{6})(?P<discard_4>ACG)
(?P<umi_1>.{8})(?P<discard_5>GAC).*" \
--extract-method=regex --stdin raw/1A_1.fastq.gz \
--stdout umitools/umi_1A_1.fastq.gz --read2-in raw/1A_2.fastq.gz \
--read2-out=umitools/umi_1A_2.fastq.gz --error-correct-cell \
--filter-cell-barcode --whitelist=whitelist/1A_whitelist.txt > log/1A_umi_extract.log;
```

#### Trimming of reads

Trimming of adaptor sequences (ILLUMINACLIP:~/Adapters/ddSeq_SureCell.fa:2:30:10)
Removal of low quality reads (LEADING:3 TRAILING:3 SLIDINGWINDOW:4:15)
Remove reads shorted than 36bp (MINLEN:36)

```
#Trimmomatic v0.36
TrimmomaticSE -threads 10 -phred33 umitools/umi_1A_2.fastq.gz trimmomatic/TRIM_1A_2.fastq \
ILLUMINACLIP:~/Adapters/ddSeq_SureCell.fa:2:30:10 \
LEADING:3 TRAILING:3 SLIDINGWINDOW:4:15 MINLEN:36 2> log/1A_trim.log;
```

#### Alignment of reads

```
#STAR v 2.7.1a
~/STAR-2.7.1a/bin/Linux_x86_64/STAR \
--genomeDir ~/Genomes/GRCm38 \
--readFilesIn trimmomatic/TRIM_1A_2.fastq \
--runThreadN 10 --outFileNamePrefix STAR/1A/1A --outFilterMultimapNmax 1 \
--outSAMtype BAM SortedByCoordinate;
```

#### Assigment of reads to genes

```
#featureCounts v1.6.0
featureCounts -a /mnt/f/Genomes/GRCm38/gencode.vM8.annotation_with_Luc.gtf \
-o featureCounts/1A -R BAM STAR/1A/1AAligned.sortedByCoord.out.bam -T 10 -s 1;
```

#### Sorting and indexing of BAM file

```
#samtools 1.7
#Using htslib 1.7-2
```

```
samtools sort featureCounts/1AAligned.sortedByCoord.out.bam.featureCounts.bam \
-o featureCounts/1A_sorted.bam;
samtools index featureCounts/1A_sorted.bam;
```

#### Generating of final count table

```
#UMI-tools version: 1.0.0
umi_tools count \
--per-gene --gene-tag=XT --assigned-status-tag=XS --per-cell \
-I featureCounts/1A_sorted.bam -S count/1A.tsv --wide-format-cell-counts \
> log/1A_umi_count.log;
```

### ANALYSIS OF RESULTS

#### Loading all packages

```
library("SingleCellExperiment")
library("scater")
library("NormExpression")
library("DropletUtils")
library("scran")
library("ggplot2")
library("Seurat")
```

```
packageVersion("SingleCellExperiment")
```

```
## [1] '1.4.1'
```

```
packageVersion("scater")
```

```
## [1] '1.10.1'
```

```
packageVersion("NormExpression")
```

```
## [1] '0.1.0'
```

```
packageVersion("DropletUtils")
```

```
## [1] '1.2.2'
```

```
packageVersion("ggplot2")
```

```
## [1] '3.1.1'
```

```
packageVersion("Seurat")
```

```
## [1] '3.0.2'
```

#### List of samples

```
##   SampleName temperature inhibitor Replicate Index_number Index_sequence
## 1         1A        37°C        no         A         N701       TAAGGCGA
## 2         2A        37°C       yes         A         N702       CGTACTAG
## 3         3A         4°C        no         A         N703       AGGCAGAA
## 4         4A         4°C       yes         A         N704       TCCTGAGC
## 5         1B        37°C        no         B         N705       GGACTCCT
## 6         2B        37°C       yes         B         N706       TAGGCATG
## 7         3B         4°C        no         B         N707       CTCTCTAG
## 8         4B         4°C       yes         B         N718       GGAGCTAC
```

#### Creating SingleCellExperiment object

```r
for(i in 1:nrow(samples)){
  sample<-as.vector(samples$SampleName[i])
  count_tsv_path <- paste0("./count/",sample,".tsv")
  count <- read.csv(count_tsv_path,header = TRUE,sep="\t")

  colnames(count)[-1] <- paste0(colnames(count)[-1],"_",samples$SampleName[i])

  annotation <- data.frame(cell=colnames(count),
                           temperature=rep(samples$temperature[i],ncol(count)),
                           inhibitor=rep(samples$inhibitor[i],ncol(count)),
                           replicate=rep(samples$Replicate[i],ncol(count)),
                           sample=rep(samples$SampleName[i],ncol(count)),
                           condition=rep(paste0(samples$temperature[i],":",
                                                samples$inhibitor[i]),ncol(count)),
                           row.names = colnames(count))
  all_count <- merge(all_count,count, by="gene", all = TRUE)
  all_count[is.na(all_count)] <- 0
  all_annotation <- rbind(all_annotation,annotation)
}


#Load all data into SingleCellExperiment object
sceseq <- SingleCellExperiment(assays = list(counts = as.matrix(all_count)),
                               colData = all_annotation)
#Remove undetected genes
keep_features <- rowSums(counts(sceseq) > 0, na.rm = TRUE) > 0
sceseq <- sceseq[keep_features,]

#Calculate empty drops
e.out = emptyDrops(counts(sceseq))
#Filter out: empty drops and cells wich less then 200 or more then 5.000 UMIs
is.cell = (e.out$FDR <= 0.05)
w2kp = which(is.cell & e.out$Total >= 200 & e.out$Total <= 5000)
sceseq = sceseq[,w2kp]
```

#### Seurat analysis

```r
# Change SingleCellExperiment object into Seurat object
seurat <- as.Seurat(sceseq, counts = "counts", data = NULL)
# Normalization of the data
seurat <- NormalizeData(seurat, verbose = FALSE)
seurat <- FindVariableFeatures(seurat, selection.methods = "vst",nfeatures = 4000)
# Run the standard workflow for visualization and clustering
seurat <- ScaleData(seurat, verbose = FALSE)
seurat <- RunPCA(seurat, npcs = 30, verbose = FALSE)
# t-SNE and Clustering
seurat <- RunTSNE(seurat, reduction = "pca", dims = 1:30, verbose = FALSE)
seurat <- RunUMAP(seurat, reduction = "pca", dims = 1:30, verbose = FALSE)
seurat <- FindNeighbors(seurat, reduction = "pca", verbose = FALSE)
seurat <- FindClusters(seurat, resolution = 0.5, verbose = FALSE)

#Removal of clusters identified as "debris"
seurat <- RenameIdents(seurat,'0' = "debris",'5' = "debris")
```

```
seurat.subset <- subset(seurat, idents = "debris", invert = TRUE)
```

**Reanalysis of seurat data without debris**

```
# Run the standard workflow for visualization and clustering
seurat.subset <- ScaleData(seurat.subset, verbose = FALSE)
seurat.subset <- RunPCA(seurat.subset, npcs = 30, verbose = FALSE)
# t-SNE and Clustering
seurat.subset <- RunTSNE(seurat.subset, reduction = "pca", dims = 1:30, verbose = FALSE)
seurat.subset <- RunUMAP(seurat.subset, reduction = "pca", dims = 1:30, verbose = FALSE)
seurat.subset <- FindNeighbors(seurat.subset, reduction = "pca", verbose = FALSE)
seurat.subset <- FindClusters(seurat.subset, resolution = 0.5, verbose = FALSE)

# Identification of markers of populations
markers_Tumour <- FindConservedMarkers(seurat.subset,
                                       ident.1 = "0",
                                       grouping.var = "condition",
                                       verbose = FALSE)

# Rename of the groups
seurat.subset <- RenameIdents(seurat.subset,
                              '0' = "Tumour cells",
                              '1' = "Tumour cells",
                              '2' = "Macrophages",
                              '3' = "T cells",
                              '5' = "T cells",
                              '4' = "NK cells")
```

```
sessionInfo()
```

```
## R version 3.5.2 (2018-12-20)
## Platform: x86_64-w64-mingw32/x64 (64-bit)
## Running under: Windows 10 x64 (build 18362)
##
## Matrix products: default
##
## locale:
## [1] LC_COLLATE=Slovak_Slovakia.1250  LC_CTYPE=Slovak_Slovakia.1250
## [3] LC_MONETARY=Slovak_Slovakia.1250 LC_NUMERIC=C
## [5] LC_TIME=Slovak_Slovakia.1250
##
## attached base packages:
## [1] parallel  stats4    stats     graphics  grDevices utils     datasets
## [8] methods   base
##
## other attached packages:
##  [1] Seurat_3.0.2                scran_1.10.2
##  [3] DropletUtils_1.2.2          NormExpression_0.1.0
##  [5] scater_1.10.1               ggplot2_3.1.1
##  [7] SingleCellExperiment_1.4.1  SummarizedExperiment_1.12.0
##  [9] DelayedArray_0.8.0          BiocParallel_1.16.6
## [11] matrixStats_0.55.0          Biobase_2.42.0
## [13] GenomicRanges_1.34.0        GenomeInfoDb_1.18.2
## [15] IRanges_2.16.0              S4Vectors_0.20.1
## [17] BiocGenerics_0.28.0
```

```
##
## loaded via a namespace (and not attached):
##   [1] Rtsne_0.15               ggbeeswarm_0.6.0
##   [3] colorspace_1.4-1         ggridges_0.5.1
##   [5] dynamicTreeCut_1.63-1    XVector_0.22.0
##   [7] BiocNeighbors_1.0.0      listenv_0.7.0
##   [9] npsurv_0.4-0             ggrepel_0.8.1
##  [11] codetools_0.2-16         splines_3.5.2
##  [13] R.methodsS3_1.7.1        lsei_1.2-0
##  [15] knitr_1.22               jsonlite_1.6
##  [17] ica_1.0-2                cluster_2.0.7-1
##  [19] png_0.1-7                R.oo_1.22.0
##  [21] sctransform_0.2.0        HDF5Array_1.10.1
##  [23] httr_1.4.0               compiler_3.5.2
##  [25] assertthat_0.2.1         Matrix_1.2-16
##  [27] lazyeval_0.2.2           limma_3.38.3
##  [29] htmltools_0.3.6          tools_3.5.2
##  [31] rsvd_1.0.1               igraph_1.2.4
##  [33] gtable_0.3.0             glue_1.3.1
##  [35] GenomeInfoDbData_1.2.0   RANN_2.6.1
##  [37] reshape2_1.4.3           dplyr_0.8.3
##  [39] Rcpp_1.0.1               gdata_2.18.0
##  [41] ape_5.3                  nlme_3.1-137
##  [43] DelayedMatrixStats_1.4.0 gbRd_0.4-11
##  [45] lmtest_0.9-36            xfun_0.6
##  [47] stringr_1.4.0            globals_0.12.4
##  [49] irlba_2.3.3              gtools_3.8.1
##  [51] statmod_1.4.30           future_1.13.0
##  [53] edgeR_3.24.3             zlibbioc_1.28.0
##  [55] MASS_7.3-51.1            zoo_1.8-5
##  [57] scales_1.0.0             rhdf5_2.26.2
##  [59] RColorBrewer_1.1-2       yaml_2.2.0
##  [61] reticulate_1.11.1        pbapply_1.4-0
##  [63] gridExtra_2.3            stringi_1.4.3
##  [65] caTools_1.17.1.2         bibtex_0.4.2
##  [67] Rdpack_0.10-1            SDMTools_1.1-221
##  [69] rlang_0.4.0              pkgconfig_2.0.2
##  [71] bitops_1.0-6             evaluate_0.13
##  [73] lattice_0.20-38          ROCR_1.0-7
##  [75] purrr_0.3.2              Rhdf5lib_1.4.3
##  [77] htmlwidgets_1.3          cowplot_0.9.4
##  [79] tidyselect_0.2.5         plyr_1.8.4
##  [81] magrittr_1.5             R6_2.4.0
##  [83] gplots_3.0.1.1           pillar_1.3.1
##  [85] withr_2.1.2              fitdistrplus_1.0-14
##  [87] survival_2.43-3          RCurl_1.95-4.12
##  [89] tsne_0.1-3               tibble_2.1.1
##  [91] future.apply_1.3.0       crayon_1.3.4
##  [93] KernSmooth_2.23-15       plotly_4.8.0
##  [95] rmarkdown_1.12           viridis_0.5.1
##  [97] locfit_1.5-9.1           grid_3.5.2
##  [99] data.table_1.12.2        metap_1.1
## [101] digest_0.6.18            tidyr_0.8.3
## [103] R.utils_2.8.0            munsell_0.5.0
```

```
## [105] beeswarm_0.2.3           viridisLite_0.3.0
## [107] vipor_0.4.5
```
